# Geno2pheno[bNAbs]: Interpretable and accurate HIV antibody resistance prediction

**DOI:** 10.1101/2025.09.11.675561

**Authors:** Martin Pirkl, Philipp Schommers, Micha Böhm, Joachim Büch, Rolf Kaiser, Thomas Lengauer

## Abstract

**Background:** Antiretroviral therapy (ART) is a life saving option for people living with HIV-1 (PLWH) and is effective against many viral strains. The most common ARTs involve combinations of drugs targeting viral or cellular proteins. Most of these drugs have to be taken daily. An alternative to ARTs with established inhibitors comprises broadly neutralizing antibodies (bNAbs). However, bNAbs share the problem of viral resistance with protein inhibitors. We developed a web service geno2pheno[bNAbs] that allows users to upload viral genotypes and estimates the respective resistance to many common bNAbs. The service uses trained statistical models to classify the virus into sensitive and resistant, respectively or to regress the IC50.

**Methods:** We used two linear models as well as two neural nets for each task and multi-task (MT) learning to train both models for IC50 prediction and classification simultaneously. During multi-task learning we penalize divergence of class and IC50 score in addition to the loss individual to each of the models.

**Findings:** We compared the linear models of geno2pheno[bNAbs] to other state-of-the-art methods like recurrent neural nets and self-attention, and found them to be competitive in regard to accuracy and have the benefit of fast computation and being easily interpretable in regard to features, i.e., positions on the envelope.

**Interpretation:** We developed a web service for the prediction of antibody resistance (geno2pheno[bNAbs]) to HIV-1, which is free to use and can be extended to other viruses, like Sars-Cov2, in the future.

## 1 Introduction

HIV continues to pose major health challenges with many deaths, especially in countries with poor access to therapeutic options [1, 2]. Antiretroviral therapy (ART) is a useful option for people living with HIV (PLWH) and is effective against many viral strains [3]. However, most ARTs constitute still a relatively large inconvenience for PLWH. The most common ARTs involve combinations of drugs targeting viral or cellular proteins. Most of these drugs have to be taken daily [4, 5]. An alternative to ARTs with protein inhibitors comprises broadly neutralizing antibodies (bNAbs). These have been shown to yield long-term viral suppression and have to be administered only once every several weeks or even months. However, bNAbs share the problem of viral resistance with protein inhibitors [6].

Solutions that are accurate, interpretable and easy to use are necessary to tackle the problem of predicting viral resistance to bNAbs. Preliminary work showed that this is possible depending on the statistical method and the individual bNAb [7, 8, 9, 10]. We developed the web service geno2pheno[bNAbs] that allows users to upload one or more viral genotypes and uses trained models to predict resistance to many common bNAbs. The predictions consist of classification (sensitive or resistant) and the regression of the half maximal inhibitory concentration (IC50).

We inspected several different models for prediction. We used support vector machines (SVM) with a linear kernel for classification and regression [11, 12]. These models are trained independently and can incur a large discrepancy between predicted class probabilities and regression values, e.g., large predicted IC50 values but simultaneously a low probability of resistance, which is contradictory. As an alternative we used a framework that allows us to train a logistic and a linear regression model (LIN) jointly. This multi-task (MT) approach allows us to penalize such contradictory predictions of the two models during training. In addition this can provide a regularization to prevent overfitting to the training data. Other multi-tasking models we consider are recurrent neural networks (RNN) with a gated recurrent unit (GRU, [13, 14, 15]) and self-attention (SA) [16]. The training and test data was downloaded from the CATNAP database [17].

Linear models are preferable due to their inherent interpretability and due to their speed of prediction. Both are favorable for a web service covering a large set of bNAbs and addressing a broad user group. The neural networks are far more complex and not necessarily more accurate than the linear models and incur far larger prediction times.

We developed the web service for antibody resistance (geno2pheno[bNAbs]), which is free to use and can be extended to other viruses, like Sars-Cov2, in the future.

## 2 Materials and Methods

### 2.1 Data processing

We downloaded virus sequences and IC50 scores for virus antibody pairs from the CATNAP data base [17] (https://www.hiv.lanl.gov/components/sequence/HIV/neutralization/). We aligned every sequence to the envelope region (ENV) of a reference sequence.

The nucleotide sequences used in the analysis were aligned to the HIV reference AF033819 from https://www.ncbi.nlm.nih.gov/nuccore/AF033819 with the tool ‘mafft’ [18]. The aligned sequences include nucleic acid substitutions and deletions. Insertions were not explicitly included. Positions where insertions were detected were marked by an asterisk as a model feature to indicate an arbitrary insertion of any length and any pattern. E.g., if we detected nucleic acid A at position 100 and then an insertion of length 10 at position 101 followed by an G at 111, position 100 was converted to A* and position 101 was set to G. The aligned sequences were converted to machine-readable format by one-hot encoding [19] (Figure 1). Each position was extended to 22 features. Twenty features represent the 20 amino acids and two features represent insertions and deletions. E.g., the wild-type virus at position 1 was presented as a vector of 21 zeros and a one for amino acid feature M (Figure 1). The full ENV sequence with a length of 857 amino acids was therefore extended to 18854 features.

**Figure 1.**
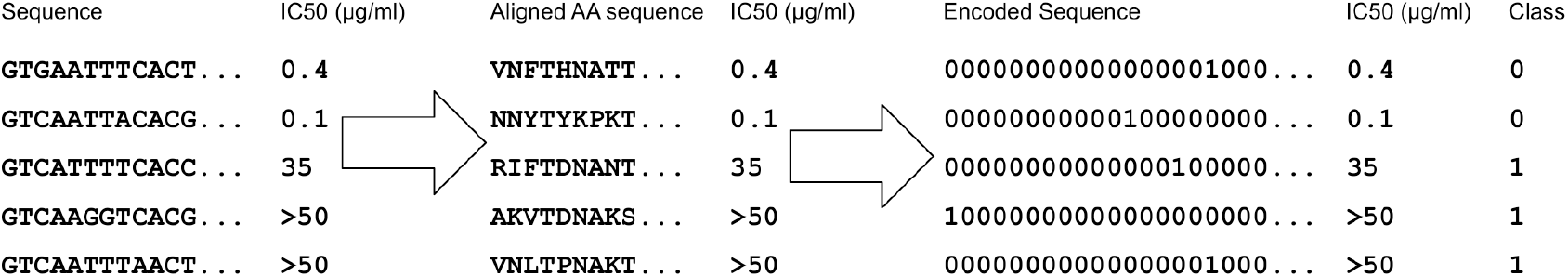
Raw sequence processing. Nucleic acid sequences are aligned and transformed to amino acid sequences. At each position the amino acid is extended to a binary sequence. The IC50 values are binarized based on a cutoff. Reproduced from [6].

The IC50 was left unprocessed in its original real-valued format. Depending on the assay and source of the available IC50 values, the values can be left- and right-censored with different lower and upper limits in μg/ml. Censored values were set to the limit of detection, e.g., >50 was set to 50. For classification, we binarized the IC50 to 0 (sensitive) and 1 (resistant) with a cutoff of 10μg/ml. We chose this cutoff according to the intermediate sensitivity classification of the Los Alamos CATNAP tool (https://www.hiv.lanl.gov/components/sequence/HIV/neutralization/main.comp). Classification has the advantage of removing noise from the continuous values. E.g., it is not important if the real IC50 is 32 or 10 because both values fall into class 1 (above 10, resistant). The disadvantage is that we lose information because a prediction of class 0 does not give further information on the real IC50 value between 0 and 10μg/ml.

We adjusted for the problem of class imbalance by up-sampling. I.e., we randomly selected with replacement from the smaller class. We avoided data leakage by making sure that all replicates from one sample group either in the training set or in the test set.

### 2.2 Choice of bNAbs

We chose 57 bNAbs to include in our web service based on two criteria (Table 1). We included bNAbs, for which enough samples were available, i.e., bNAbs, which had at least 100 sensitive and 100 resistant samples available after binarization of the IC50. However, we also included bNAbs with a smaller sample size, if they were of interest to the field of research.

**Table 1:**
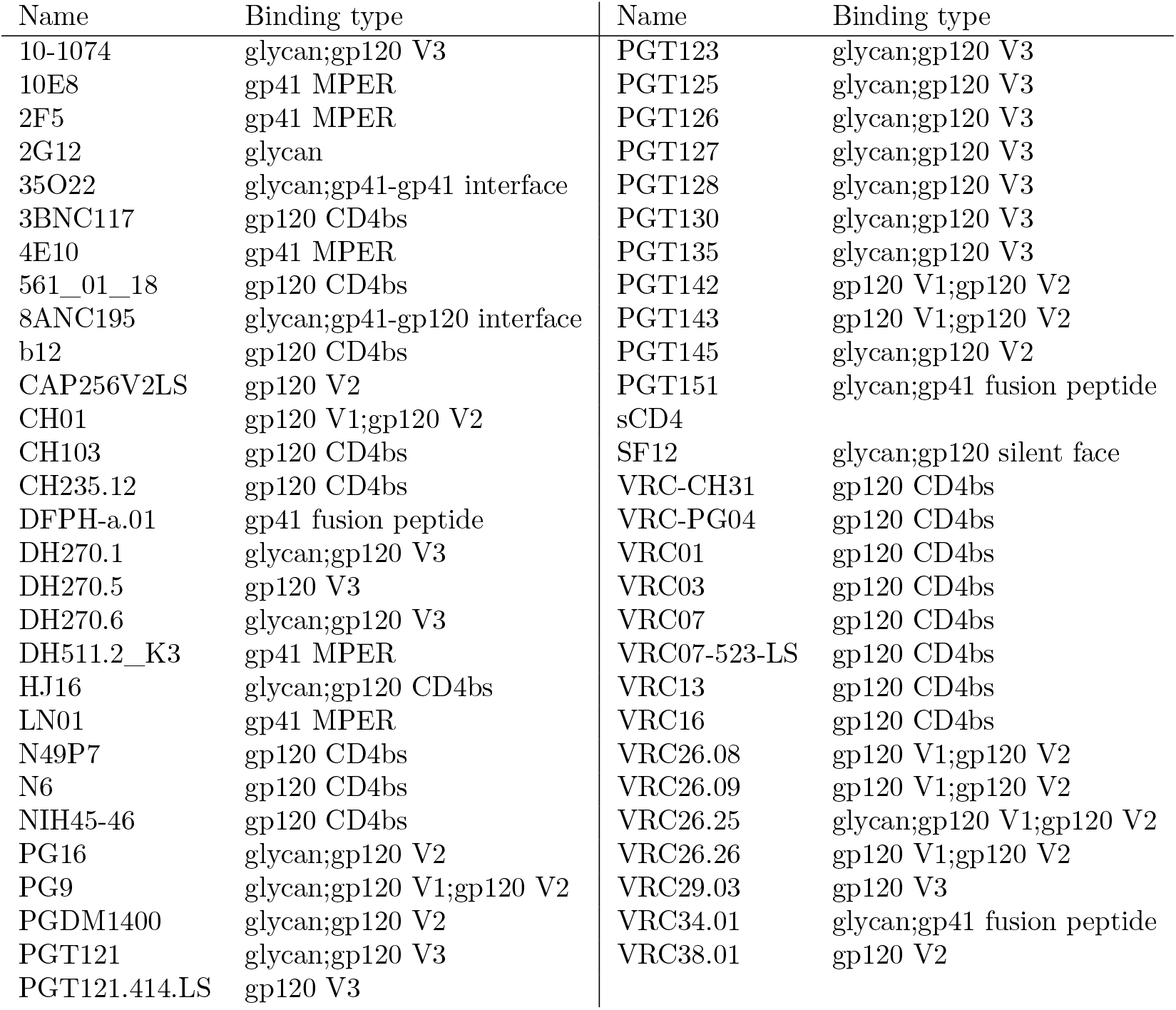
BNAbs available in our prediction system. Binding types were extracted from the CAT-NAP database [17].

### 2.3 Models

Our base line model was the well established state-of-the-art method of support vector machines (SVM) for C-classification and for *ϵ*-regression with linear kernels [11]. Cost *C* ∈ {2^*−i*^} and *ϵ* ∈ {10^*i*^} with *i* ∈ 1, 3, 5, 7, 9, 11 are hyperparameters that were optimized during ten-fold cross validation repeated five times. The cost factor determines the trade-off between maximizing the margin (class separation) and minimizing the classification error. The *ϵ* for regression defines how much a regressed point is allowed to deviate from the regression line without penalty. The classification and regression models were trained independently with the Julia [20] package ‘LIBSVM.jl’ [21, 22, 23].

The models competing with the SVM were all trained with multi-tasking. For the three multitask models, linear/logistic regression (LIN), a Gated Recurrent Unit (GRU) and Multihead Self-attention (SA), we used the Julia package ‘FLUX.jl’ [24, 25]. The models were optimized by gradient descent minimizing a loss function. Linear and logistic regression were simple linear transformations of all feature values to an scalar output, which is applied to the logistic function. Each pairing of position and amino acid is a feature. Features without information, e.g., zero in all samples, were removed. The RNN consisted of a single GRU layer ([13],v3) with 22 input and 22 output features fed back into the input layer of the next position. The GRU propagated through all 857 positions of the ENV sequence twice, once from position 1 to position 857 and after a reset, forgetting the previous positions, back from position 857 to position 1. The final output values for each position and each direction were transformed in the same fashion as for the linear and logistic regression models (Figure 2). The multihead self-attention model (SA) uses multiple attention heads the number of which is an additional hyper-parameter. The SA model is applied to all 857 positions each with 22 input and 22 output features. We keep model complexity low by using the number of heads as the hidden attention layer. The output was transformed in the same fashion as in the previous models. We could only remove features in the GRU and SA models, if a whole position was uninformative, e.g., exhibits the exact same binary pattern over the amino acids for all samples. For this reason, the data for the GRU and SA models is less sparse than for the linear models in general. But the GRU and SA models are computationally more expensive due to two additional reasons. First, both models use more computations than the linear models when applied to a sequence. Second, the data has to be transformed to explicitly include each sequence position (Figure 3). The linear models (LIN,SVM) are unaware of the actual sequence positions. Overall, we trained eight different models, four for classification and four for regression. In the context of this manuscript we will be referring to the four methods used (SVM, LIN, GRU, SA) and only distinguish between classification and regression in detailed descriptions. This is not a significant restriction, because the classification models differ from the regression models only by the use of a sigmoid function applied to the output. All four models are listed in Table 2.

**Table 2:** The four models trained for resistance prediction. The top two rows show the four models and their abbreviations used in this manuscript. The other rows indicate whether classification and regression was trained jointly (multi-task, MT) and what hyperparameters needed optimization.

| Model Abbrev. | Multihead self-attention<br>SA | Gated Recurrent Unit<br>GRU | Logistic/linear regression<br>LIN | Support vector machine<br>SVM |
| --- | --- | --- | --- | --- |
| multi-task | ✓ | ✓ | ✓ | × |
| Cost value | × | × | × | ✓ |
| $\epsilon$ | × | × | × | ✓ |
| Epochs | ✓ | ✓ | ✓ | × |
| Batchsize | ✓ | ✓ | ✓ | × |
| Dropout % | ✓ | ✓ | ✓ | × |
| N heads | ✓ | × | × | × |

**Figure 2.**
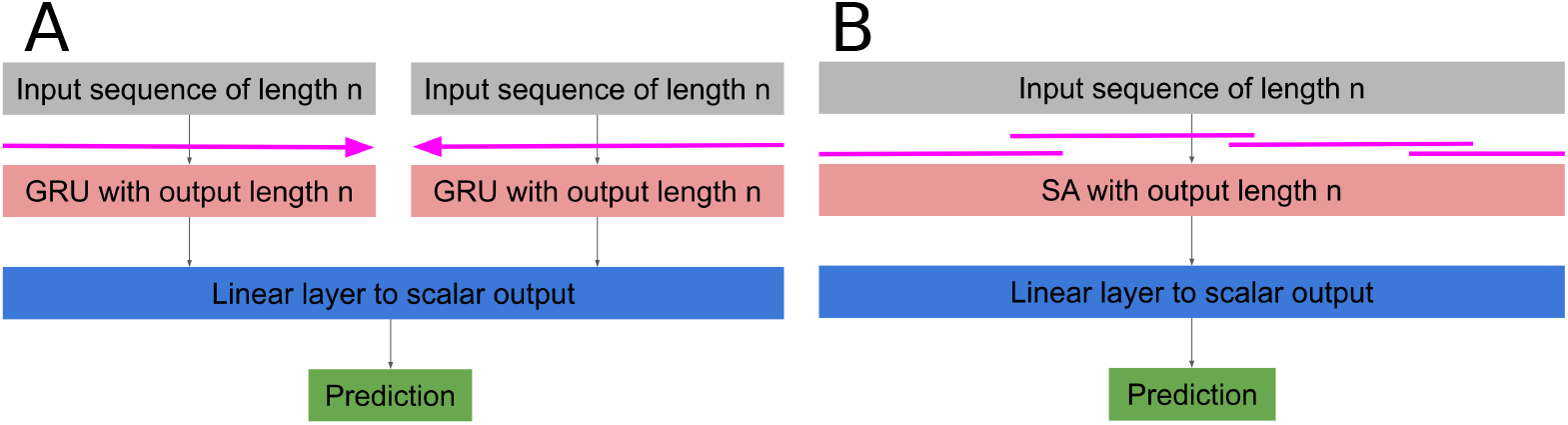
multi-task models. SA uses the individual sequence as input and the output is projected linearly to a scalar. The attention heads focusing on different regions of the sequence are visualized above the attention layer. The third layer of the SA model is the same layer type used for the LIN model. In the model layout of the GRU (B) the sequence is independently traversed in both directions (horizontal arrows). The output of both sequence results is jointly linearly projected to a scalar. The logistic function is applied to the prediction in case of classification.

**Figure 3.**
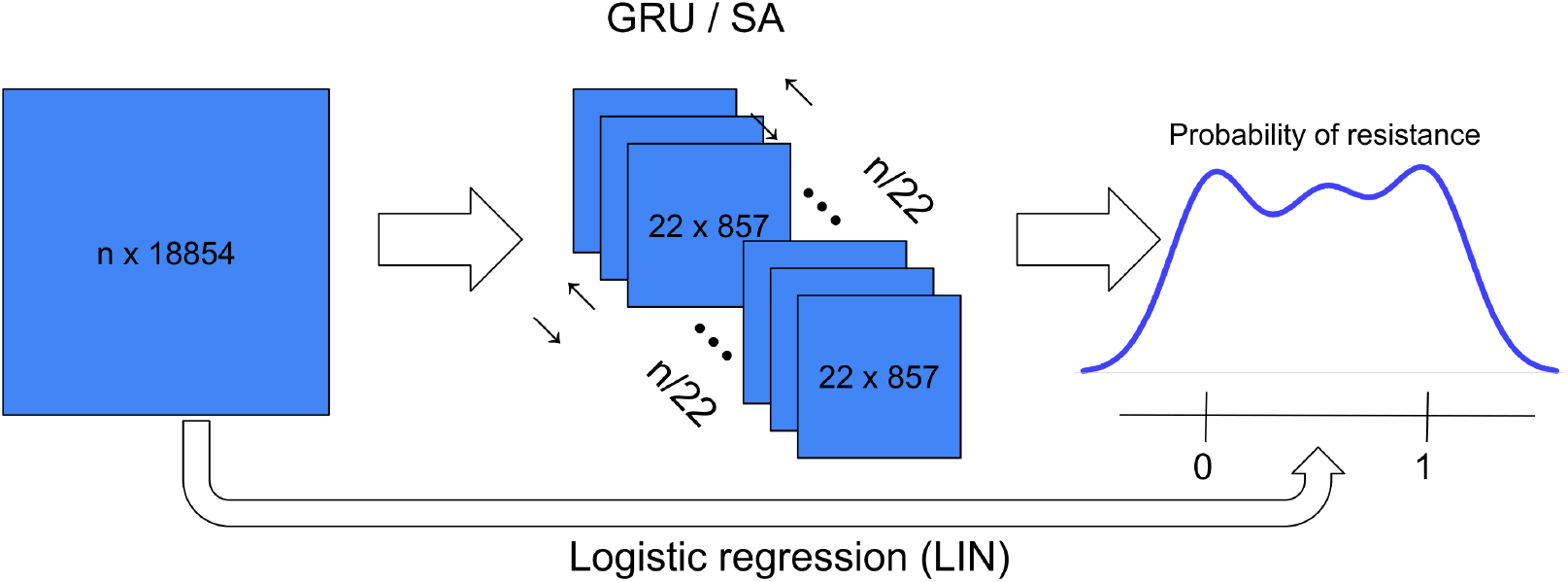
Data projection including all features. The logistic regression model transformed the input data of *n* samples directly with a linear projection into *n* probabilities. The *n* × 18854 input matrix had to be transformed to a 22 × 857 × *n* array for the GRU and SA models. Each sample was represented by a sequence of length 857 and 22 features at each position.

#### Hyperparameters

The hyperparameters (Table 2) for the three multi-task models (LIN, GRU, SA) were the number of epochs (training iterations) ∈ {10, 50, 100}, the batch size (number of samples to process before a descent) ∈ {8, 16, 32}, the percentage of dropout [26] ∈ {0, 25, 50} and for the SA model the number of heads ∈ {1, 4, 8}. The percentage of dropout randomly sets a percentage of edges in the model to 0 (removes them) for regularization purposes.

#### Loss functions

For the three multi-task models we add binary cross-entropy [27], Eq. 1 for class probabilities, huber loss (*δ* = 1, [28], Eq. 2) for the predicted IC50 and the negative covariance (*Cov*) of both Eq. 3, such that, if the covariance of predicted class probabilities and predicted IC50 is large, the loss is small. The explicit formulation of the three loss functions are

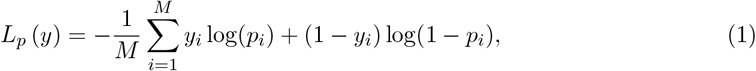

with the classification label *y*_*i*_ ∈ {0, 1} (sensitive, resistant) and the predicted probability of resistance *p*_*i*_ ∈ [0, 1] for *i* ∈ {1, …, *M*}.

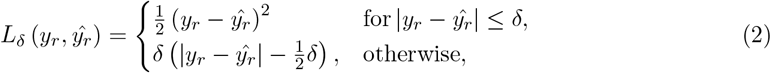

with *y*_*r*_ as the real IC50 value and *ŷ*_*r*_ as the predicted IC50 value. Both losses are combined with the covariance for the multi-task loss in

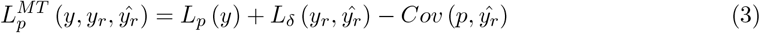

with *p* = (*p*_1_, *…, p*_*M*_). Since all models except the SVM are trained with multi-tasking, we only use Eq. 3.

### 2.4 Split into training set and test set

We randomly split the data into a training set and a test set with a ratio of nine to one. We trained the hyperparameters with ten-fold cross validation on the training set. We repeated the ten-fold cross validation five times with differently sampled folds. We ensured that all replicates of a sample were grouped in a single fold. We trained the final model on the full training set with the optimal hyper-parameter setting.

We assessed the accuracy during cross validation with the area-under-receiver operating characteristic (AUC) for the classification task and with the mean squared error (MSE) for the regression task. Previous work suggests the AUC as a reasonable performance indicator for balanced data [7, 8, 9, 10].

The final model was validated on the test set with the AUC on the classification task and Harrell’s concordance index (HCI,[29]) on the regression task. We used the MSE during training and the HCI for testing, because the MSE forces the regression model to predict values at the correct magnitude. The HCI is scale invariant and is more interpretable, because it ranges from 0 to 1 with an expected value of 0.5 for random predictions.

### 2.5 Converting class probability to discrete classes

In our web service we provide a prediction of either sensitive, intermediate resistance and full resistance. For this we discretized the predicted class probability based on cutoffs determined from our training data.

We picked 100 equidistant points within the interval of the maximum and minimum of predicted raw values, not yet converted to probabilities, of the training set. For two of those non-overlapping points we discretized the predictions into three classes. We used the AUC to score the discretization. We greedily searched through all unique, non-overlapping pairs of points to find the optimal cutoff. We used 100 bootstrap runs to estimate an optimal cutoff distribution. We picked the median of each distribution as our final cutoffs. This process is illustrated in Figure 4. The distribution of sensitive and resistant samples are shown in green and red, respectively. The cutoffs for intermediate and full resistance prediction are the yellow and blue lines, respectively. The estimated distribution for both cutoffs are colored correspondingly.

**Figure 4.**
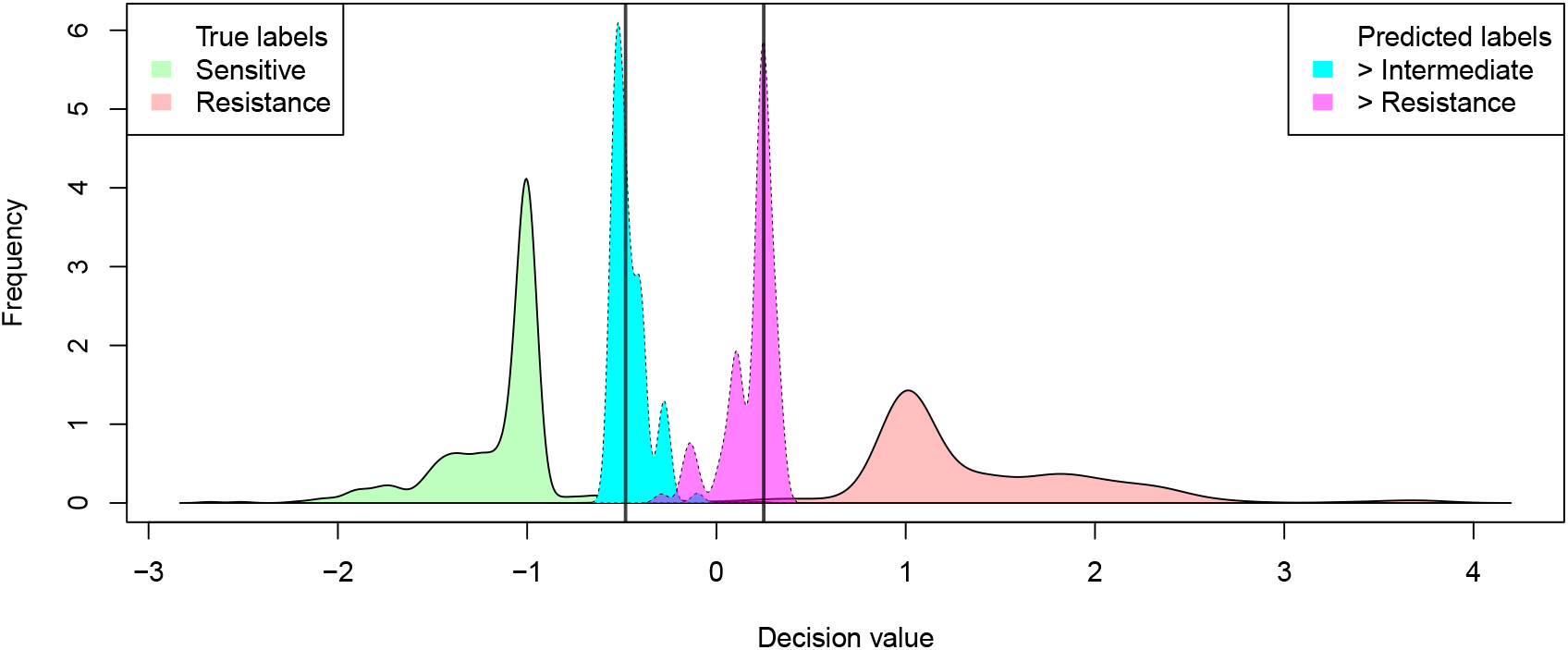
Example from real data for estimated cutoffs based on greedy search combined with bootstrapping. The green distribution depicts the sensitive samples and the red distribution the resistant samples. The bootstrapped distributions for the two cutoffs are shown in light blue, from sensitive to intermediate resistance, and in purple, from intermediate resistance to resistance. The medians of both bootstrapped distributions are shown as black vertical lines and chosen as the final cutoffs.

## 3 Results

### 3.1 Training accuracy from cross validation

The GRU model showed a significantly higher accuracy for the classification than the other models (Figure 5, p-value*<* 0.01 for Wilcoxon rank sum test). The SA model is significantly worse (p-value<0.001). The distribution of the accuracy for the regression were all virtually identical.

**Figure 5.**
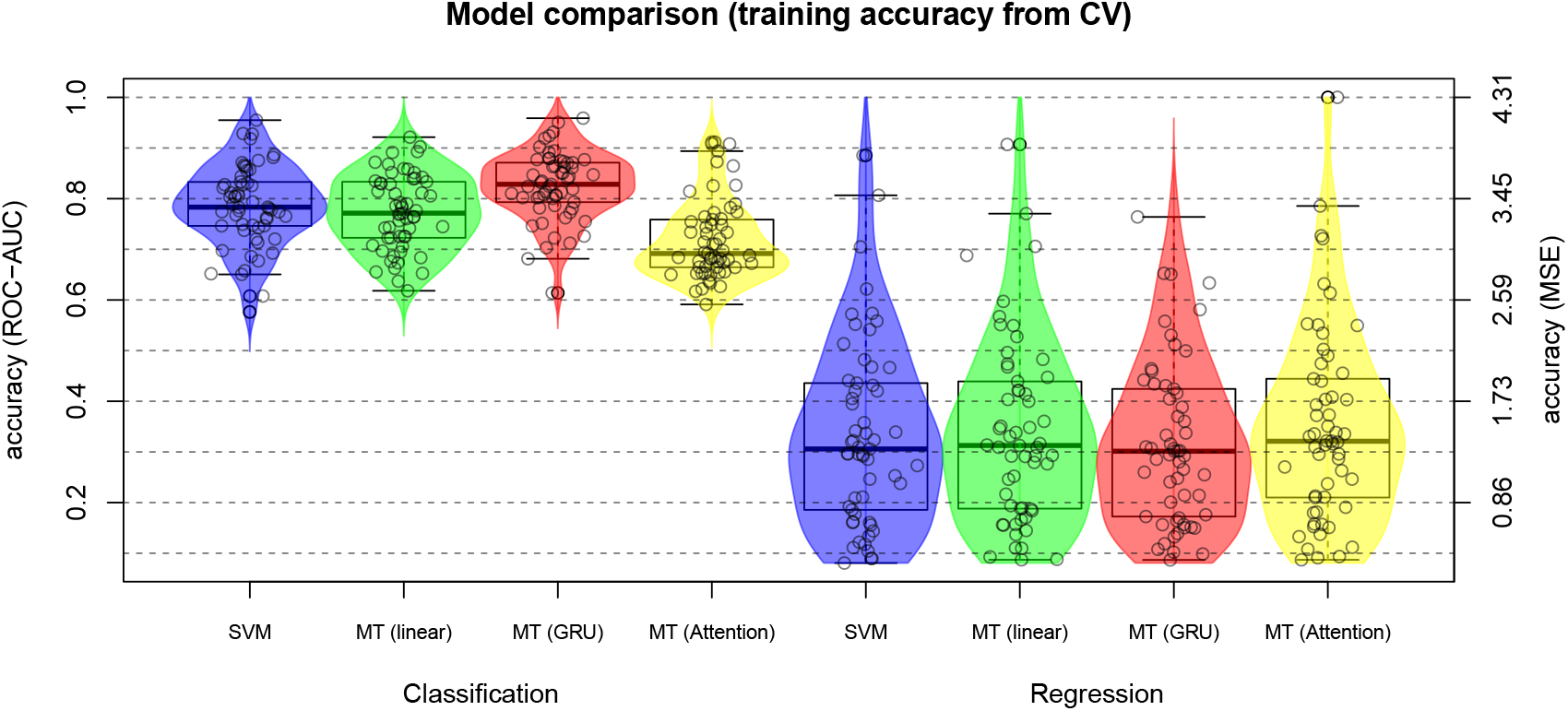
Accuracy of the four classification and regression models.

We analyzed whether the training accuracy is correlated with the number of samples for each bNAb. We did so with a simple linear regression of the accuracy depending on the number of samples. For all models there was a significant positive correlation for classification accuracy (p-values<0.01 and <0.05 for the GRU) and significant negative correlation with the regression MSE (p-values<0.05). This is expected because, in general, we would expect the AUC to be higher and the MSE to be lower the more samples are available, i.e., the models become more accurate the more samples become available. The effect for classification was most significant for the SA model.

We also stratified the training accuracies by bNAb binding type (Table 1, Wilcoxon rank sum test). There was no significant difference in classification accuracy per binding type. The MSE for regression accuracy was significantly worse (greater) in the group of bNAbs with binding type gp120 V1/2 (p-value<0.05, median in [0.55, 0.55, 0.51, 0.56]) than in the other binding type groups. For bNAbs in group gp41 MPER the MSE was significantly smaller (p-value<0.01, median in [0.13, 0.14, 0.13, 0.14]) than for the other binding types. The models showed no significant difference in the regression accuracies in other bNAb binding type groups.

We also looked for enrichment of ENV contact and binding positions of each bNAb for both linear models (SVM, LIN). ENV contact and binding positions were downloaded from the Los Alamos HIV Sequence Database (https://www.hiv.lanl.gov/). This information was not available for all bNAbs. A Wilcoxon rank sum test calculated 4 (SVM classification), 20 (MT classification), 6 (SVM regression) and 14 (MT regression) bNAbs models out of 38 each as significantly enriched (FDR<0.1, Benjamin-Hochberg correction) in the respective bNAbs contacts and binding position on the ENV. However, we observed an inherent position-wise symmetry in each position with 22 possible features each (Figure 6). Summing up all weights virtually equals 0. I.e., even though the linear models do not explicitly encode the 857 positions, the one-hot encoding already has that positional information implied by virtually having only a single one (sometimes more due to uncertainty about several amino acids for a position) and 21 zeros for every 22 positional features. I.e., any position with large positive weights for one or more amino acids (resistance) has either also large negative weights for one or more other amino acids or many negative weights for many amino acids. Hence, the Wilcoxon rank sum test applied to all weights is expected to have little power. If we perform the test with all weights greater than zero, hence only including features favoring resistance, all positive model weights were significantly enriched with large weights at the bNAbs contact and binding positions.

**Figure 6.**
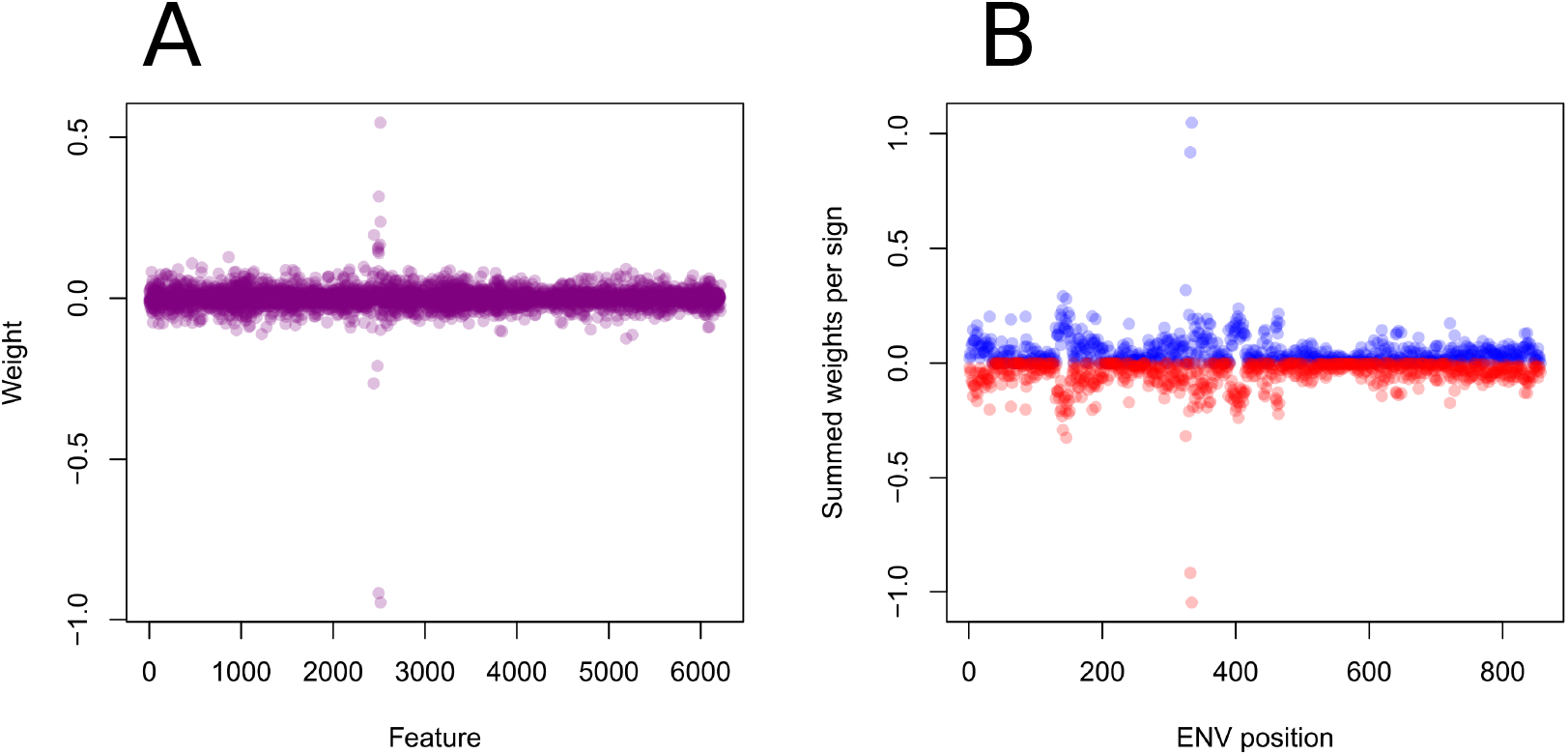
Example of model weights. The raw model weights (A) show many non-zero weights with some outliers for both signs (negative, positive). Summed up per position and sign (B), each position’s positive and negative weights are virtually equal.

**Figure 7.**
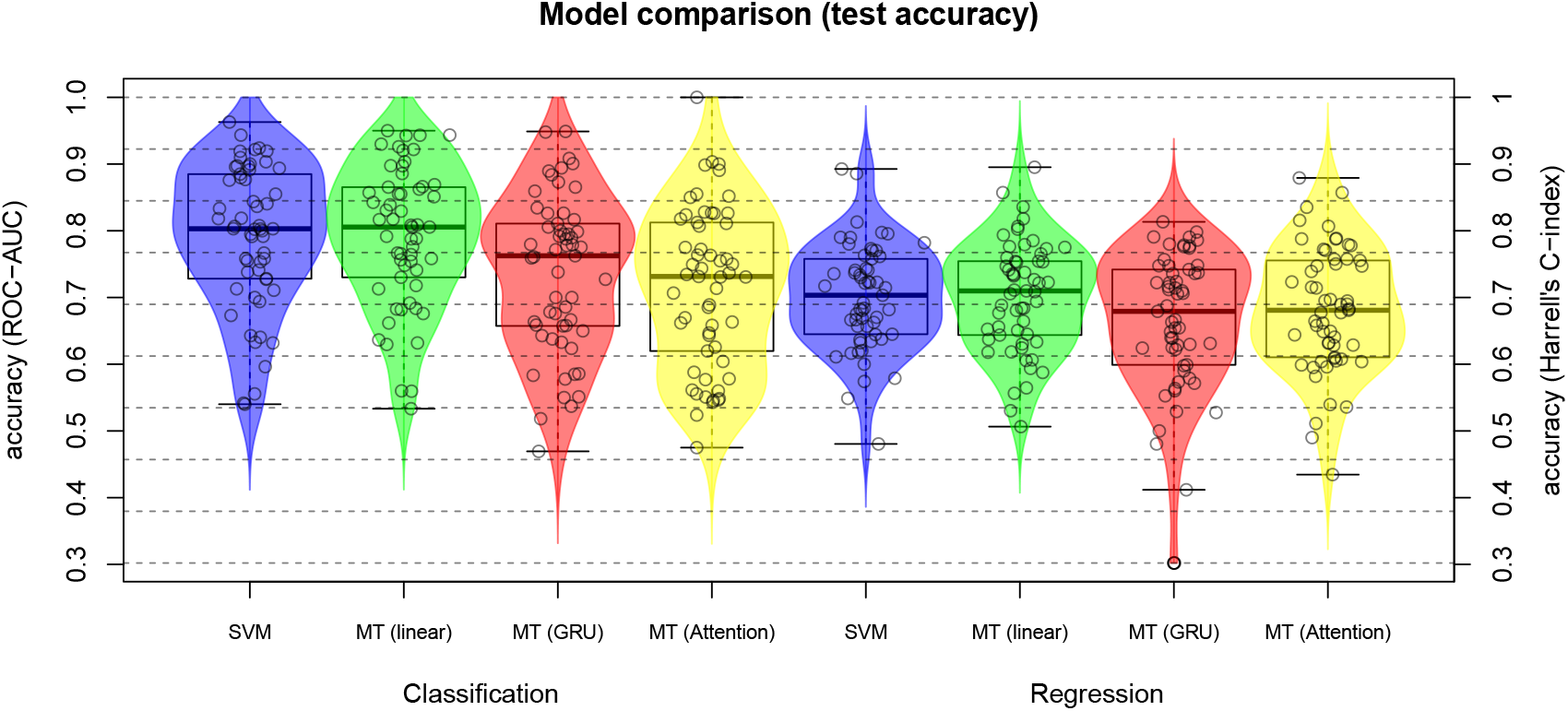
Validation accuracy. Analogous to the cross validation accuracy but on the test set not used during training of the models. The accuracy measure for the IC50 prediction is Harrell’s concordance index.

Overall (Figure 5) the two linear models seem to not perform much worse than the GRU, but are much easier to interpret and also faster, which is very important for a web service to ensure responsiveness. Hence, we concentrate most of the following analyses (sections 3.3 and 3.4) on the two linear models used for our web-service (SVM, LIN).

### 3.2 Testing accuracy

The highest scoring models over all bNAbs on the test set were the linear models with GRU at third and SA at last rank. As before, all models did equally well on the regression task. For the regression we used the MSE to train the models, but Harrell’s concordance index for better interpretation. All models were significantly above a coin flip approach with a theoretical accuracy of 0.5 for both measures.

### 3.3 Multi-tasking correlates classification and regression

As described above, we used multi-task learning for logistic and linear regression (LIN). The goal was to avoid the prediction of a low probability of resistance simultaneously with a high IC50 value. This can generally happen, if both models were trained independently. We reason that training both models jointly should mitigate that negative effect. While multi-task learning can be viewed as a method for regularization, it does not guarantee any increase in model accuracy as for any regularization method. In the following sections we compare the independently trained SVM models for classification and regression with the jointly trained linear multi-task models (LIN).

Correlation of class probability and IC50 prediction over all training samples was significantly higher in the multi-task model (LIN, Figure 8, A, p-value*<* 0.01 for Wilcoxon signed rank test). I.e., the multi-ask model is more likely than the independent model to predict a higher class probability for the same sample, if it predicts a larger IC50 and vice versa. Especially the bNAbs with a correlation between IC50 and class probability over all samples of below 0.3 were at least at correlation 0.4 or higher. The greatest increase was achieved for bNAb HJ16 from a correlation between IC50 and class probability over all samples of 0.23 for the SVM to 0.88 for the multi-task model (LIN). However, for some bNAbs (21) the correlation decreased. The decrease was mostly very small or at an already high level between 0.7 and 1. The greatest decrease was observed for 3BNC117 from a solid correlation of 0.59 down to 0.43. If the independently trained model has a higher training accuracy for IC50 or class prediction, it is chosen over the multi-task model.

**Figure 8.**
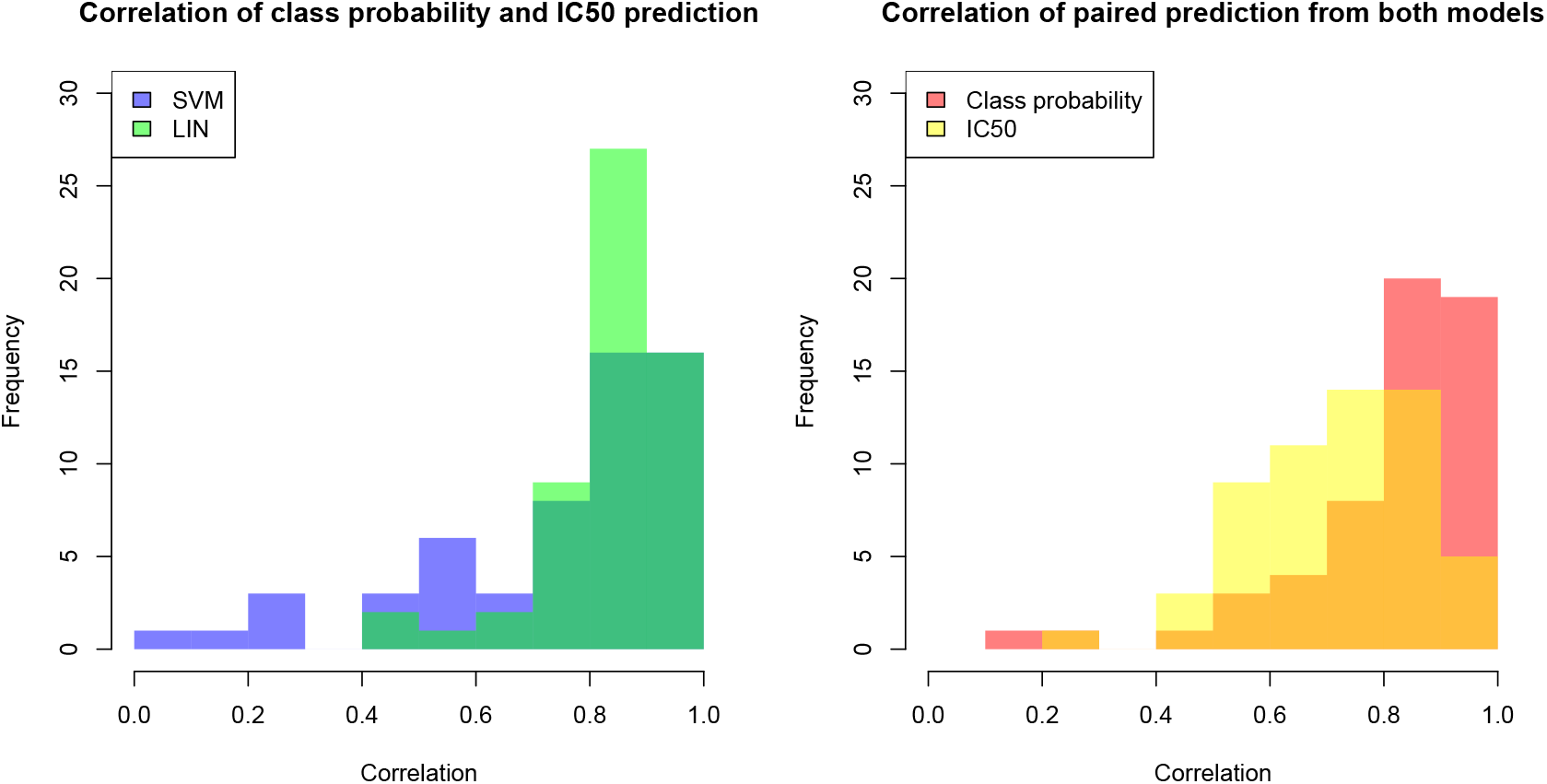
Correlation of model predictions. Correlation within models (A) and correlation across models (B). The correlation of prediction within a model shows how well predicted class probability correlates with IC50 for the same method. Correlation across models shows how well predictions (either class or IC50) correlated between the two models (SVM and multi-task).

Just because class probability and IC50 prediction correlate within an individual model, does not mean that the predictions correlate across models. Figure 8 (B) shows that predictions by both models, SVM and LIN, correlated relatively highly for both prediction types, class probability and IC50. Especially the predictions for class probability reach a very high agreement between both models except for the outliers N6 (0.14) and 3BNC117 (0.24). Almost perfect agreement for the class probability was reached for 2F5 (0.98). The picture is similar for IC50 prediction although the correlation between the two models is a bit lower, in general. HJ16, which exhibited the highest increase in correlation of predicted class probability and IC50 due to multi-task learning, had the lowest agreement between IC50 prediction from the SVM and the multi-task model. The highest agreement between the model predictions was achieved by PGT143 (0.93). We can also observe a general trend that if the independently trained SVM models for a bNAb had highly correlated predictions of IC50 and class probability, i.e., the multi-task learning was not necessary, we also observed high agreement across model predictions between the SVM and the multi-task model and vice versa, e.g., for bNAb HJ16.

We inspected the correlation of predicted class probability and IC50 in more detail. The example in Figure 9, A shows one bNAb, which seemed to have benefited highly from multi-task learning. Contrary to the SVM prediction (blue), the multi-task predictions (green) show virtually no samples for which class probability and IC50 contradict each other. The example in Figure 9, B shows an example that showed no improvement. The IC50 predictions from the multi-task model were much more spread out compared to the SVM and produced actually more contradicting predictions. The example in Figure 9, C on one hand shows improvements regarding samples with high class probabilities, while samples with low class probabilities showed the same artifact as the example in Figure 9, B.

**Figure 9.**
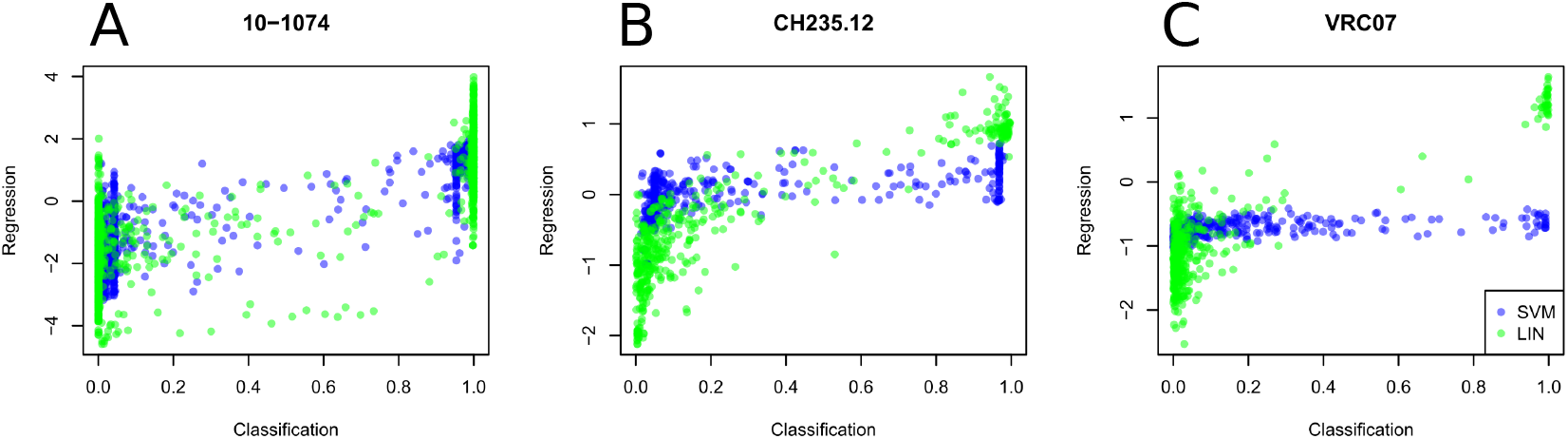
Three examples for the correlation between predicted class probabilities and IC50. Predicted was the full data set including test samples by the linear SVM, independently trained, and the multi-task model (LIN). A: The multi-task models fails to correlates some data pairs. B: The multi-task model successfully correlates the predicted class probability with the predicted IC50 values. C: The multi-task models achieves high agreement for predicted class probabilities close to 1 and corresponding predicted IC50 values, but predicts an almost random IC50 distribution for class probabilities close to 0.

### 3.4 The web service

For the final model for each bNAb and each prediction type (class probability, IC50) we choose between the SVM and the LIN model based on the cross validation accuracy from model training. The final models are free to use at our web service https://bnabs.geno2pheno.org/. The backend is relatively lightweight, because the linear models were implemented as a vector multiplication.

Data in form of a consensus sequence can either be pasted in via a form (ADD SEQUENCE button) or uploaded in fasta file format, allowing multiple sequences at once (UPLOAD FASTA FILE button). The uploaded sequences are aligned against a reference genome and translated into amino acids sequences. No user data, like sequence information, is in any way stored on our server.

After pressing the ANALYZE button, the user is transferred to the PREDICTION tab (Figure 10). On this tab the user can select any of the sequences uploaded via fasta. The predictions show the alphabetically listed bNAbs and some statistics like the raw decision value used to compute probabilities and the Z-score putting the prediction in context with the training data. Next to the prediction, class probability or IC50, the top 15 positions and their respective amino acids are shown ranked by their absolute weight in each model and colored for favoring sensitive (green) or resistance (red), respectively. The colored symbols indicate sensitive (green), intermediate resistance (yellow) and full resistance (red) of the sequence to the respective bNAb. The colors are based on cutoffs estimated from the training data.

**Figure 10.**
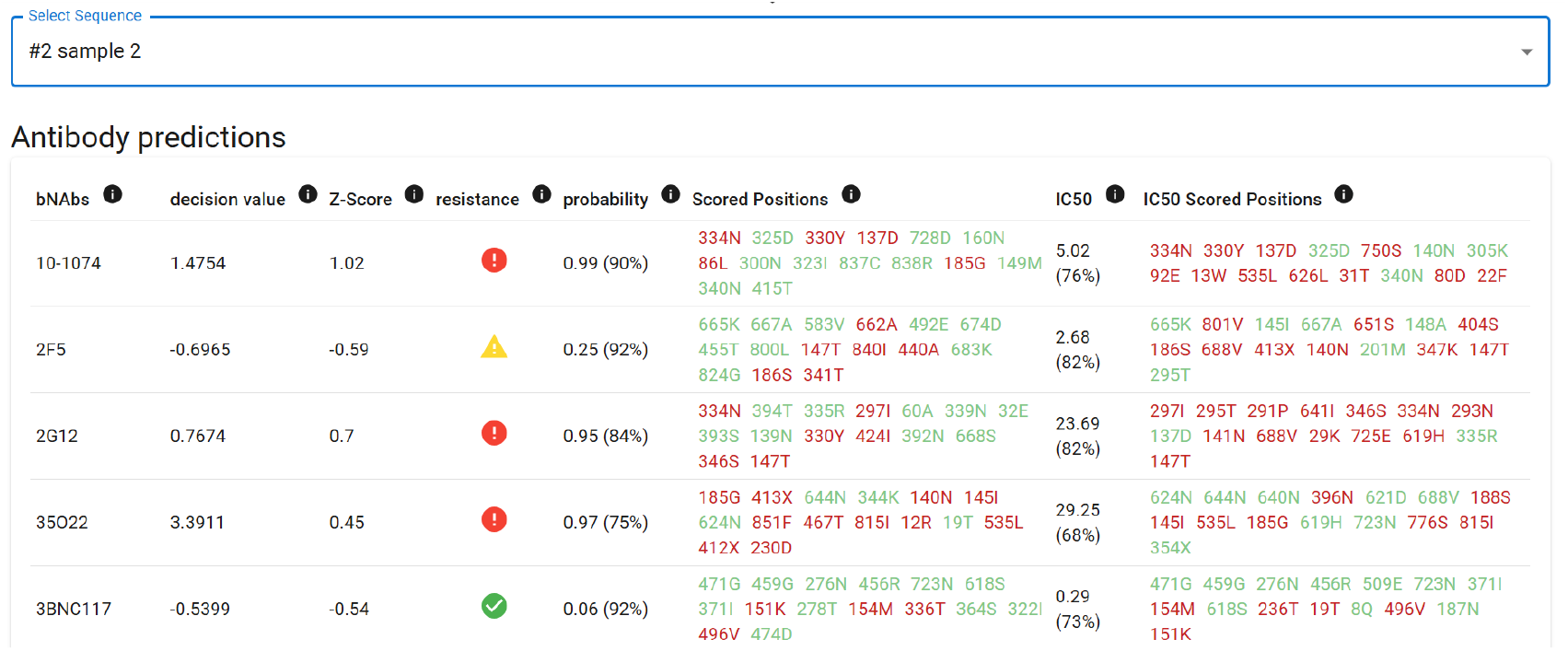
PREDICTION tab of our geno2pheno[bNAbs] web service. All previously analyzed bNAbs are ordered alphabetically. Shown are statistics (decision value, Z-score) and the resistance levels based on our cut offs in green, yellow and red, from sensitive to intermediate to full resistance. Additionally to the predicted probability and IC50 we show the confidence of our prediction for both models in parentheses. We also show the 15 positions on the analyzed sequence with largest absolute weights.

The web service also includes the prediction of the HIV-1 subtype and shows the alignment to the HIV-1 reference.

## 4 Discussion

We have introduced a free to use, speedy web service for the prediction of resistance to 57 different bNAbs. During the training of the linear models (SVM, LIN) implemented in the web service’s backend, we compared them to other state-of-the-art machine learning methods. Some of the competing models indicated higher predictive performance for the class prediction (Figure 5, accuracy median (classification): GRU (82.8%), SVM (78.3%)). However, the complexity and training effort do not seem to be worth the increase, especially in the context of novel incoming data and updating the models. While the linear models take minutes to hours of training, the more complex models take from days to months of cpu time. It should also be noted that the linear models come with an inherent way of interpreting the results, while other methods would need non-trivial additional complexity to provide interpretation of the results [30].

BNAb binding site type does not seem to make any difference for classification. However, gp41 MPER were significantly more accurate for IC50 prediction, while gp120 V1/2 bNAbs were predicted significantly less accurately by all models. Interestingly, while the competing models show partially significantly higher accuracy during training, they show lower accuracy for the test set not used for training. This is of course a problem, because if we were not biased on selecting the linear models due to their stated benefits, we would select the best model according to the cross validation accuracy. In many cases our model choice would be the recurrent neural network with a gated recurrent unit. This model, however, does not seem to generalize well to unseen data, but this could also be due to the random selection of the test data.

Enrichment analysis showed that for many bNAbs informative linear model weights, i.e., those with large values, were significantly enriched in corresponding bNAb contact and binding position on the ENV. I.e., even though the models use all possible combinations of position and amino acids as features, biologically relevant features for resistance are included to a significant number. This indicates that our models partially detect inherent biological processes. It also explains why rules based models based on these processes, i.e., mutations, already achieve accuracies significantly greater than random guessing [31].

Another issue we addressed is the independence of the two models, one for classification and one for regression (IC50). In general the classification task is easier, i.e., reaches higher accuracy. While we lose information during binarization to class labels (0, 1), we also remove noise. E.g., our classification model treats a IC50 value of 3 and 8 the same and ignores their difference. The regression models try to predict this difference. To tackle this problem of the IC50 regression, we used multi-task learning. That means we trained both models jointly by putting a penalty (negative covariance) on divergence of predicted class probability and IC50. If the class probability is high and the IC50 low or vice versa the loss increases (equation 3). Another benefit of this method could be that it may act as a regularization mechanism. We put stronger constraints on the model during training, which may lead to better generalization on predicting data not used for training. As a caveat, multi-task learning does not necessarily improve accuracy. I.e., even if our predictions appear to be better, because both models are aligned, the independently trained models could be more accurate. This is indeed the case, because the SVM gets chosen over the multi-task model (LIN) for most bNAbs (37 out of 57 for classification and 31 out of 57 for regression).

Another limitation of our web service is that the models were trained from freely available data. This data is not necessarily from a single source, i.e., different assays could have been used to produce IC50 values. This is an additional factor of noise and bias and could be a disadvantage to our approach. However, many bNAbs reached a class prediction performance (AUC) of greater than 80 or even 90%, which seems to indicate that this issue might not be too relevant.

We also included a small number of bNAbs of interest to the field, which had only a small set of samples available, e.g., due to novelty of the bNAb. This made the training and validation unstable, i.e., very large variance of accuracy. Test accuracy might be skewed due to the small number of test samples.

In conclusion, we implemented a free and easy to use web service geno2pheno[bNAbs] https://bnabs.geno2pheno.org/ with a high prediction accuracy for many bNAbs of interest. In the future this web service can be updated when new data is available or new research of bNAbs can be implemented. This web service can also be extended to other viruses.

## Data availability

All data used is available from the Los Alamos National Laboratory at https://www.hiv.lanl.gov/components/sequence/HIV/neutralization/.

## Acknowledgement

This research has been conducted in the context of the Viromarkers project (grant agreement No 101194735).

We furthermore thank the Regional Computing Center of the University of Cologne (RRZK) for providing computing time on the DFG-funded (Funding number: INST 216/512/1FUGG) High Performance Computing (HPC) systems CHEOPS and RAMSES as well as support.

## Author contributions

MP, PS, RK and TL conceptualized the study. MP conducted the statistical analysis under supervision of TL. MP implemented the models into the web service. MP, MB and JB build the web service application. MP, PS, RK and TL interpreted the results. MP, RK and TL wrote the initial draft of the manuscript. All authors have read and agreed to the final draft of the manuscript.

### Conflict of interest

none declared

## Notes

### Competing Interest Statement

The authors have declared no competing interest.

### Summary of Updates

Additional acknowledgement by recognizing a European research grant.

